# Cadmium-associated differential methylation throughout the placental genome: epigenome-wide association study of two US birth cohorts

**DOI:** 10.1101/130286

**Authors:** Todd M. Everson, Tracy Punshon, Brian P. Jackson, Ke Hao, Luca Lambertini, Jia Chen, Margaret R. Karagas, Carmen J. Marsit

## Abstract

**Background:** Cadmium (Cd) is a ubiquitous toxicant that during pregnancy can impair fetal development. Cd sequesters in the placenta where it can impair placental function, impacting fetal development. We aimed to investigate Cd-associated variations in placental DNA methylation (DNAM), associations with gene expression, and identify novel pathways involved in Cd-associated reproductive toxicity.

**Methods:** Using placental DNAM and Cd concentrations in the New Hampshire Birth Cohort Study (NHBCS, n=343) and the Rhode Island Child Health Study (RICHS, n=141), we performed an EWAS between Cd and DNAM, adjusting for tissue heterogeneity using a reference-free method. Cohort-specific results were aggregated via inverse variance weighted fixed effects meta-analysis, and variably methylated CpGs were associated with gene expression. We then performed functional enrichment analysis and tests for associations between gene expression and birth metrics.

**Results:** We identified 17 Cd-associated differentially methylated CpG sites with meta-analysis p-values < 1e-05, two of which were within a 5% false discovery rate (FDR). Methylation levels at 9 of the 17 loci were associated with increased expression of 6 genes (5% FDR): *TNFAIP2*, *EXOC3L4*, *GAS7*, *SREBF1*, *ACOT7*, and *RORA*. Higher placental expression of *TNFAIP2* and *ACOT7*, and lower expression of *RORA*, were associated with lower birth weight z-scores (p-values < 0.05).

**Conclusion:** Cd associated differential DNAM and corresponding DNAM-expression associations at these loci are involved in inflammatory signaling and cell growth. The expression levels of genes involved in inflammatory signaling (*TNFAIP2*, *ACOT7*, and *RORA*), were also associated with birth metrics, suggesting a role for inflammatory processes in Cd-associated reproductive toxicity.

**Significance:** Cadmium is a toxic environmental pollutant that can impair fetal development. The mechanisms underlying this toxicity are unclear, though disrupted placental functions could play an important role. In this study we examined associations between cadmium concentrations and DNA methylation throughout the placental genome, across two US birth cohorts. We observed cadmium-associated differential methylation, and corresponding methylation-expression associations at genes involved in cellular growth processes and/or immune and inflammatory signaling. This study provides supporting evidence that disrupted placental epigenetic regulation of cellular growth and immune/inflammatory signaling could play a role in cadmium associated reproductive toxicity in human pregnancies.

## Background

Cadmium (Cd) is a toxic heavy metal that is released into the environment from mining and industrial processes, the application of phosphate fertilizers, and fossil fuel combustion; it accumulates in soils and is readily taken up by plants (1). Human exposure primarily occurs via consumption of contaminated foods, or through using tobacco products (2). Cadmium serves no biological role in humans, and high concentrations can cause severe health consequences such as kidney damage, bone and joint problems and various cancers (1). However, trace exposures to cadmium may also have toxic effects, particularly to pregnant mothers and the developing fetus (3).

Recent epidemiologic studies have observed associations between Cd concentrations in maternal and/or fetal tissues with restricted fetal growth and pregnancy complications (4–10). The toxic mechanisms that underlie these associations are not clear, though Cd is known to induce oxidative stress, interfere with cell-cycle regulation, and alter apoptotic signaling at the cellular level (11). Additionally, Cd tends to sequester in the placenta throughout pregnancy, limiting the direct fetal exposure, though its accumulation within the placenta is likely not without consequence (12). Animal models and *in vitro* studies have suggested that maternal Cd exposure during pregnancy can perturb maternal-fetal nutrient and waste transfer (13, 14), increase oxidative stress in placental tissue (15), interfere with placental glucocorticoid synthesis (16), and alter trophoblast proliferation and apoptosis (17). Thus, the placenta appears to be a critical target tissue for the toxic effects of Cd during human pregnancies.

We hypothesize that various Cd-associated perturbations to placental functions are likely linked to the selective expression or repression of specific biological pathways. Likewise, maternal Cd exposure during pregnancy has been associated with variations in DNA methylation (DNAM), an epigenetic mechanism that regulates gene expression potential, in maternal and/or fetal blood samples (18). To date, only one study has investigated associations between placental Cd and DNAM throughout the placental genome in humans. Mohanty et al (2015) explored Cd-associated DNAM in a small study of 24 placentae, identifying multiple differentially methylated loci nearby or within genes involved in cell-damage, angiogenesis, cell-differentiation, and organ development; these associations differed by fetal sex (19). Studies with larger samples using recent methodological advances to estimate and adjust for tissue heterogeneity are needed to understand how Cd exposure influences the placental epigenome of human pregnancies, and whether such variations are related to fetal growth restriction.

We conducted a study of the associations between placental Cd and placental DNAM throughout the epigenome across two well characterized independent cohorts while examining the potential impact of tissue heterogeneity on these associations. Secondarily, we investigated whether the top hits from our analysis were enriched for biological pathways consistent with findings from other studies, whether differentially methylated loci were associated with the expression levels of nearby genes, and whether these variations in expression were associated with standardized measures of birth weight, birth length and head circumference.

## Results

The EWAS for this study included 343 mother-infant pairs drawn from the New Hampshire Birth Cohort (NHBCS) and 141 mother-infant pairs from the Rhode Island Child Health Study (RICHS). The most notable differences between the NHBCS and RICHS samples (Table 1) were with racial diversity and birth size. Because RICHS was over-sampled for LGA and SGA infants by design, we observed a higher proportion of SGA and LGA infants compared to NHBCS. Placental samples from the RICHS cohort also had slightly higher median Cd concentrations (4.34 ng/g) compared to NHBCS (3.13 ng/g), and concentrations were approximately log-normal distributed in both cohorts (Supplemental Figure 1).

**Table 1:** Maternal and offspring characteristics for the NHBCS (n=343) and the RICHS (n=141) samples.

| **Maternal Characteristics** |  | **NHBCS<sup>\*</sup>** | **RICHs<sup>\*</sup>** |
| --- | --- | --- | --- |
| Smoking During Pregnancy | Any | 21 (6.1%) | 19 (13.5%) |
|  | None | 322 (93.9%) | 122 (86.5%) |
| Educational Attainment | High-School or less | 37 (10.8%) | 34 (24.1%) |
|  | More than High-School | 306 (89.2%) | 107 (75.9%) |
| Ethnicity | Not White | 6 (1.7%) | 36 (25.5%) |
|  | White | 337 (98.3%) | 107 (74.5%) |
| **Infant Characteristics** |  |  |  |
| Gestational Age (weeks) |  | 38.95 (1.54) | 39.33 (1.00) |
| Birth Weight (grams) |  | 3435.14 (510.95) | 3623.09 (716.03) |
| Size for Gestational Age |  |  |  |
|  | Small (SGA) | 15 (4.4%) | 21 (14.9%) |
|  | Adequate (AGA) | 292 (86.1%) | 68 (48.2%) |
|  | Large (LGA) | 32 (9.4%) | 52 (36.9%) |
| Sex | Female | 159 (46.4%) | 68 (48.2%) |
|  | Male | 184 (53.6%) | 73 (51.8%) |
| **Placental Cd (ng/g)<sup>†</sup>** |  |  |  |
|  | Overall | 3.13 (2.61) | 4.37 (2.71) |
|  | Below cohort-specific median | 2.07 (1.05) | 2.78 (1.35) |
|  | Above cohort-specific median | 4.68 (2.32) | 5.49 (2.44) |

<sup>\*</sup> Means and standard deviations are presented for continuous variables, whereas frequencies and percentages are presented for categorical variables.
<sup>†</sup> Medians and IQRs presented due to skewed distributions of Cd concentrations.

We regressed DNA methylation M-values (at 408,367 CpGs in NHBCS and 397,040 CpGs in RICHS) on dichotomized placental Cd concentrations (median split) while adjusting for fetal sex, maternal smoking during pregnancy, and maternal education level, in each cohort. We then combined parameter estimates and p-values via inverse variance weighted fixed effects meta-analysis (390,583 CpGs overlapped between the cohorts) which yielded three loci with Cd-associations at a 5% FDR and 18 additional loci with p-value < 1e-05 (Table 2), in which Cd was predominantly associated with hypermethylation. Full results from the meta-analysis are provided in the supplemental materials (Supplemental Table 1).

**Table 2:** Results from cohort-specific and meta-analyses of Cd-associated DNA methylation, for probes with meta-analysis p-values < 1e-05; adjusted for maternal smoking during pregnancy, maternal education and fetal sex.

| CpG Annotations |  |  | Meta-Analysis |  |  | NHBCS (n=343) |  | RICHS (n=141) |  |
| --- | --- | --- | --- | --- | --- | --- | --- | --- | --- |
| CpG ID | Ch. | Gene | $\beta_1$ | p-value | FDR | $\beta_1$ | p-value | $\beta_1$ | p-value |
| cg16034168 | 1 | *ACOT7* | 0.39 | 1.92E-06 | 0.107 | 0.48 | 4.47E-05 | 0.31 | 7.39E-03 |
| cg18384460 | 1 | - | 0.35 | 2.22E-06 | 0.107 | 0.43 | 1.44E-05 | 0.25 | 2.42E-02 |
| cg08705382 | 1 | *GALNT2* | 0.34 | 9.51E-06 | 0.151 | 0.34 | 8.06E-04 | 0.35 | 3.79E-03 |
| cg26262055 | 4 | - | 0.22 | 8.94E-06 | 0.151 | 0.29 | 1.15E-02 | 0.21 | 2.12E-04 |
| cg26653990 | 5 | - | 0.23 | 8.75E-06 | 0.151 | 0.25 | 1.41E-04 | 0.19 | 1.87E-02 |
| cg00766220 | 5 | - | 0.14 | 4.37E-06 | 0.147 | 0.13 | 3.32E-04 | 0.14 | 4.13E-03 |
| cg09391066 | 6 | - | 0.19 | 6.10E-06 | 0.151 | 0.21 | 1.10E-03 | 0.18 | 1.65E-03 |
| cg03797906 | 7 | *TRIL* | 0.17 | 9.62E-06 | 0.151 | 0.21 | 2.48E-04 | 0.14 | 7.97E-03 |
| cg11157440 | 8 | *CSGALNACT1* | 0.29 | 7.64E-06 | 0.151 | 0.28 | 6.19E-04 | 0.30 | 3.91E-03 |
| cg21437157 | 8 | - | 0.23 | 4.12E-06 | 0.147 | 0.19 | 1.53E-03 | 0.33 | 3.32E-04 |
| cg05888894 | 8 | *EXT1* | 0.33 | 2.81E-07 | 0.046 | 0.33 | 5.56E-03 | 0.33 | 1.54E-05 |
| cg03509329 | 14 | *LRRC16B* | 0.15 | 7.09E-06 | 0.151 | 0.19 | 9.15E-04 | 0.12 | 1.47E-03 |
| cg15345741 | 14 | - | 0.28 | 4.10E-07 | 0.046 | 0.28 | 2.52E-04 | 0.29 | 4.64E-04 |
| cg17232357 | 15 | *SMAD6* | -0.19 | 2.16E-06 | 0.107 | -0.17 | 1.04E-02 | -0.21 | 5.85E-05 |
| cg12642651 | 17 | *GAS7* | 0.38 | 5.81E-06 | 0.151 | 0.27 | 2.78E-02 | 0.48 | 3.09E-05 |
| cg03762242 | 17 | *GAS7* | 0.26 | 3.38E-06 | 0.142 | 0.26 | 7.76E-04 | 0.26 | 1.34E-03 |
| cg16768966 | 17 | *GAS7* | 0.37 | 4.05E-07 | 0.046 | 0.34 | 1.55E-03 | 0.40 | 6.93E-05 |
| cg15285733 | 17 | *GAS7* | 0.17 | 7.32E-07 | 0.062 | 0.16 | 2.42E-03 | 0.18 | 8.78E-05 |
| cg02393525 | 17 | *NAGLU* | 0.23 | 8.90E-06 | 0.151 | 0.16 | 3.33E-02 | 0.31 | 3.13E-05 |
| cg06707369 | 19 | *TYROBP* | 0.17 | 7.49E-06 | 0.151 | 0.19 | 2.35E-04 | 0.15 | 8.89E-03 |
| cg11707084 | 19 | *CRX* | 0.23 | 7.79E-06 | 0.151 | 0.25 | 1.37E-04 | 0.19 | 1.68E-02 |

CpG: cytosine-phosphate-guanine methylation site; Ch.: chromosome; FDR: false discovery rate adjusted p-value; NHBCS: New Hampshire Birth Cohort; RICHS: Rhode Island Child Health Study.

Cell-specific DNAM is a well-recognized confounder in epigenetic epidemiology, leading many studies to estimate the proportions of constituent cell-types from the DNAM array data (20), and adjust for these proportions in their regression models. However, estimation of cell-type proportions requires access to cell-type specific methylomes for the individual cells that constitute the heterogenous tissue under study, and no reference methylomes currently exist for placental tissue. Thus we utilized a reference free method that deconvoluted the major axes of variation in DNAM, which are likely associated with tissue heterogeneity, to estimate the proportions of the putative constituent cell-types (PCTs) in our placental samples (21). Via this method, we estimated five and four PCTs in NHBCS and RICHS, respectively. We then tested whether the 10,000 CpGs with greatest variability in PCT-specific methylation were enriched for biologically relevant GO terms or KEGG pathways, and checked for consistency between the two cohorts. We observed 31.6% overlap among these 10,000 PCT-defining CpGs, and strong concordance in the biological functions of the top 100 GO terms (60% overlap) and KEGG pathways (76% overlap) across the two cohorts. The top 10 KEGG pathways from both cohorts included metabolic pathways, neuroactive ligand-receptor interaction, axon guidance, calcium signaling, olfactory transduction, cell adhesion molecules (CAMs), and cAMP signaling (Supplemental Tables 2 & 3, Supplemental Figure 2).

We then tested whether the distribution of these PCTs varied with placental Cd concentrations. Higher Cd was associated with increased proportions of PCT-1 (3.4% difference, T-test p-value = 0.010) and PCT-2 (4.7% difference, T-test p-value = 0.022) in NHBCS, while higher Cd was associated with increased proportions of PCT-2 (7.3% difference, T-test p-value = 0.047) in RICHS. Since these associations may confound the relationships between Cd and locus-specific DNAM, we then reproduced the cohort specific multivariable models and meta-analysis while also adjusting for the estimated PCT proportions. These adjustments substantially reduced inflation in NHBCS (λ=1.47 before, and λ=0.96 after adjustment), while only modestly increasing inflation in RICHS (λ=1.03 before, and λ=1.10 after adjustment). Furthermore, the PCT-adjustments reduced the proportion of small heterogeneity p-values (< 0.05) in the meta-analysis from 7.9% in the non-PCT adjusted models to 4.6% in the PCT-adjusted models. Full results from the PCT-adjusted meta-analysis are provided in the supplemental materials (Supplemental Table 4). The majority of Cd-DNAM associations from the original models were attenuated after PCT-adjustments. For instance, the parameter estimates for our top 3 hits from the original meta-analysis (cg15345741, cg16768966, and cg05888894) were reduced by 11%, 16%, and 45%, respectively, after adjustment for PCTs, demonstrating that cell mixture likely explained much of the Cd-associated variation for cg05888894, but only a modest proportion of the Cd-DNAM associations at cg15345741 and cg16768966. Additionally, the coefficients for cg15345741 and cg16768966 retained highly significant p-values (< 1e-05) after PCT adjustment, due to improved standard errors around these parameter estimates.

The PCT-adjusted top hit from NHBCS was cg20656525 on chromosome 14, downstream of *TNFAIP2* and *EXOC3L4*, while the top hit from RICHS was cg07708653 on chromosome 1, within the body of the *HPDL* gene. The PCT-adjusted meta-analysis yielded 17 CpGs with Cd-association p-values < 1e-05 (Table 3), two of which were significant at a 5% FDR (cg15345741 and cg11707084). Both of the FDR significant CpGs also had significant linear associations with log-Cd (cg15345741 estimate = 0.17, p-value = 0.00019; cg11707084 estimate = 0.16, p-value = 0.000014) via liner mixed models. The meta-analyses again revealed a clear trend towards higher methylation levels associated with higher Cd concentrations; 94% of Cd-associated loci with p-values < 1e-05 had positive beta coefficients. Of note, 41% of CpGs with suggestive p-values (< 1e-05) were on chromosome 17, five of which were within the *GAS7* gene. Given the high proportion of *GAS7* CpGs among the top hits from the meta-analysis, we explored the co-methylation structure within *GAS7* and as well as all Cd-DNAM associations within and around this gene. The strongest Cd-associated methylation sites tended to congregate around the first exons of three of the four different *GAS7* variants (-a, -b and -d) and exhibited a strong co-methylation structure as demonstrated by strong positive spearman correlation rho values (Figure 2).

**Table 3:** Results from cohort-specific and meta-analyses of Cd-associated DNA methylation, for probes with meta-analysis p-values < 1e-05; adjusted for maternal smoking during pregnancy, maternal education, fetal sex, and estimated tissue heterogeneity.

| CpG Annotations |  |  | Meta-Analysis |  |  | NHBCS (n=343) |  | RICHS (n=141) |  |
| --- | --- | --- | --- | --- | --- | --- | --- | --- | --- |
| CpG ID | CHR | Gene | $\beta_1$ | p-value | FDR | $\beta_1$ | p-value | $\beta_1$ | p-value |
| cg16034168* | 1 | *ACOT7* | 0.32 | 2.80E-06 | 0.171 | 0.32 | 3.11E-04 | 0.32 | 2.78E-03 |
| cg24000528 | 2 | *MIR10B* | 0.16 | 7.59E-06 | 0.182 | 0.18 | 1.52E-03 | 0.16 | 1.50E-03 |
| cg24696183 | 11 | *KCNQ1DN* | -0.16 | 4.31E-06 | 0.171 | -0.20 | 5.67E-04 | -0.14 | 1.71E-03 |
| cg23193177 | 11 | - | 0.24 | 4.68E-06 | 0.171 | 0.22 | 4.94E-04 | 0.27 | 2.70E-03 |
| cg15345741* | 14 | - | 0.25 | 1.49E-07 | 0.040 | 0.25 | 9.69E-05 | 0.24 | 4.22E-04 |
| cg06100161 | 15 | *RORA* | 0.20 | 1.97E-06 | 0.171 | 0.12 | 6.28E-02 | 0.28 | 1.99E-06 |
| cg03917020 | 16 | - | 0.14 | 5.52E-06 | 0.174 | 0.13 | 3.62E-04 | 0.14 | 4.85E-03 |
| cg02600679 | 16 | *FAM38A* | 0.25 | 4.92E-06 | 0.171 | 0.17 | 2.55E-02 | 0.34 | 2.02E-05 |
| cg04267691 | 17 | *GAS7* | 0.18 | 6.98E-06 | 0.178 | 0.12 | 3.09E-02 | 0.24 | 2.57E-05 |
| cg10517535 | 17 | *GAS7* | 0.31 | 6.46E-06 | 0.177 | 0.16 | 6.97E-02 | 0.59 | 3.08E-07 |
| cg03762242* | 17 | *GAS7* | 0.23 | 3.33E-06 | 0.171 | 0.20 | 2.08E-03 | 0.26 | 4.12E-04 |
| cg16768966* | 17 | *GAS7* | 0.31 | 1.92E-06 | 0.171 | 0.23 | 7.31E-03 | 0.42 | 3.12E-05 |
| cg10618704 | 17 | *GAS7* | 0.36 | 5.91E-06 | 0.174 | 0.24 | 2.03E-02 | 0.54 | 1.65E-05 |
| cg11851174 | 17 | *RAI1* | 0.19 | 3.69E-06 | 0.171 | 0.20 | 1.07E-04 | 0.18 | 1.10E-02 |
| cg19944656 | 17 | *TIMP2* | 0.18 | 2.62E-06 | 0.171 | 0.17 | 8.06E-05 | 0.18 | 9.82E-04 |
| cg11707084* | 19 | *CRX* | 0.22 | 2.09E-07 | 0.040 | 0.23 | 3.10E-05 | 0.22 | 1.93E-03 |
| cg03566291 | 21 | - | 0.23 | 9.46E-06 | 0.201 | 0.21 | 9.57E-04 | 0.29 | 2.38E-03 |

\* CpG Sites that yielded meta-analysis p-values < 1e-05 in models before and after heterogeneity adjustments.
CpG: cytosine-phosphate-guanine methylation site; CHR: chromosome; FDR: false discovery rate adjusted p-value; NHBCS: New Hampshire Birth Cohort; RICHS: Rhode Island Child Health Study.

**Figure 1:**
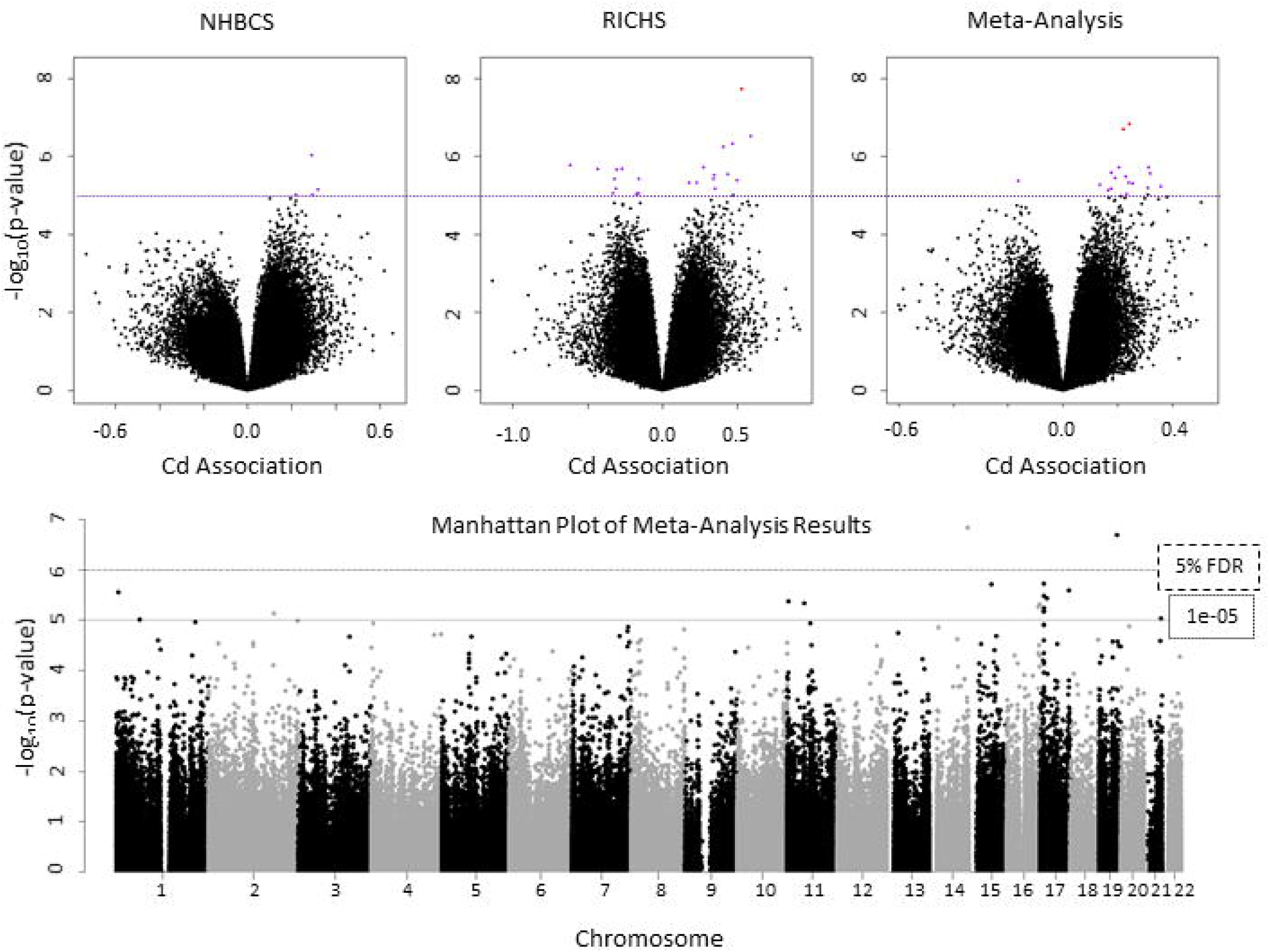
Volcano plots from each cohort-specific EWAS and the meta-analysis, and Manhattan plot of meta-analysis results; all models adjusted for fetal sex, maternal smoking during pregnancy, mother’s highest achieved education level, and putative placental cell types; dashed line represents the FDR 5% threshold, dotted line represents suggestive association (p-values < 1e-05).

**Figure 2:**
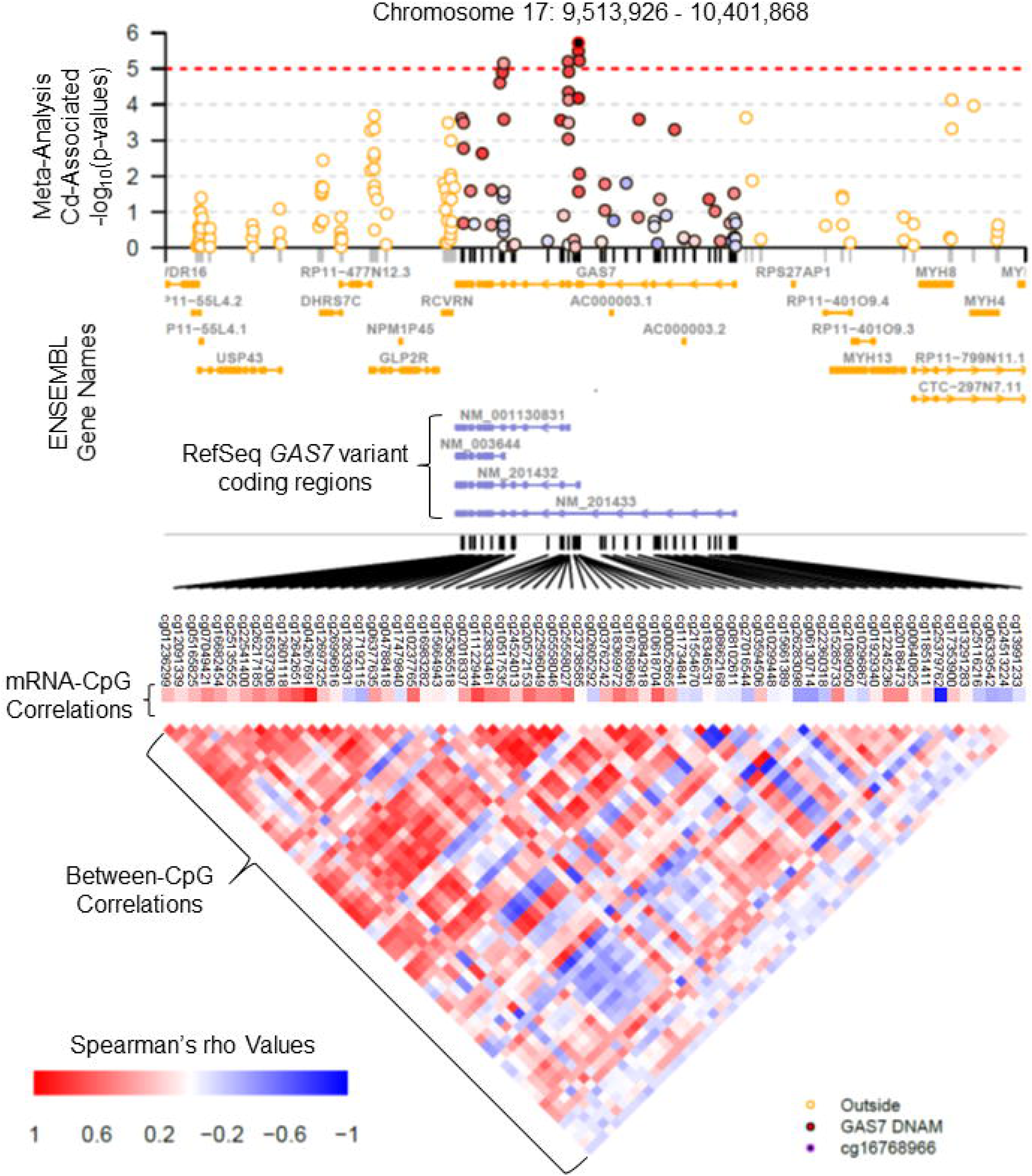
Manhattan plot of meta-analysis results surrounding the *GAS7* gene, annotated with Ensembl gene names and RefSeq IDs for *GAS7* variants, Spearman correlations between *GAS7* CpG sites, and CpG-expression correlations for *GAS7*. Cd-associated CpG outside of the *GAS7* coding region (±1,500bp), have orange circles, where CpGs within *GAS7* have red, white, or blue circles corresponding to positive, no, or negative associations, respectively.

We also investigated the underlying biology that may be affected by Cd-associated variations in the placental epigenome. Since we had too few FDR-significant findings to perform an enrichment analyses on only those hits, we tested for GO-term and KEGG pathway enrichment among the top 250 CpGs associated with placental Cd after PCT adjustment. The top GO terms included dimethylarginase activity, eye and heart valve development, and heparin sulfate proteoglycan & polysaccharide biosynthetic process (Supplemental Table 5), though none of these enrichments were significant at an FDR threshold of 5%. On the other hand, our gene set was enriched with genes from numerous KEGG pathways (Supplemental Table 6), 12 of which yielded p-values within an FDR threshold of 5% (Table 4); these included cell adhesion molecules (CAMs) and tight junctions, multiple cancer pathways, multiple G-protein and nitric oxide signaling pathways, and cytotoxic processes.

**Table 4:** Significantly enriched KEGG pathways (FDR 5%) associated with for genes annotated to the top 250 CpG sites from the PCT-adjusted meta-analysis; N represents the total genes in the KEGG path, DE represents the number of genes within our top 250 sites.

| KEGG Pathway Description | N | DE | FDR |
| --- | --- | --- | --- |
| Ras signaling pathway | 217 | 6 | 0.0058 |
| Tight junction | 163 | 5 | 0.0058 |
| Platelet activation | 122 | 4 | 0.019 |
| Cell adhesion molecules (CAMs) | 133 | 4 | 0.019 |
| Taste transduction | 79 | 3 | 0.019 |
| Apelin signaling pathway | 136 | 4 | 0.019 |
| Pancreatic cancer | 61 | 3 | 0.026 |
| Oxytocin signaling pathway | 150 | 4 | 0.026 |
| cGMP-PKG signaling pathway | 157 | 4 | 0.026 |
| Viral carcinogenesis | 191 | 4 | 0.030 |
| Metabolic pathways | 1189 | 8 | 0.034 |
| Natural killer cell mediated cytotoxicity | 107 | 3 | 0.042 |

To explore whether gene expression levels might be associated with variations in methylation levels at Cd-associated CpGs with meta-analysis p-values < 1e-05, we performed an expression quantitative trait methylation (eQTM) analysis. We regressed the expression levels of any gene within 100kb of a candidate CpG site on the methylation levels of that CpG. Three of these CpGs (cg24000528, cg03917020, and cg23193177) were not within 100kb of any genes that yielded detectable RNA from our placental samples, and thus 14 CpGs and 32 genes were included in this analysis; only *GAS7* was cis to multiple CpGs within our set of candidates. We produced 36 linear models ranging between 1 and 7 genes tested per CpG (Table 5). Of the 36 eQTM models, 10 yielded FDR-significant linear associations between methylation and expression of six unique genes (Supplemental Figure 3). Of particular note, were the associations between our top hit from the meta-analysis and the expression levels of *EXOC3L4* (β_1_ = 1.95, p-value = 0.0043) and *TNFAIP2* (β_1_ = 1.88, p-value = 0.0055). Additionally, all five of the Cd-associated *GAS7* CpGs (within or nearby the first exons of variants-a, -b, and -d) were associated with increased GAS7 expression (Table 5). On the other hand, DNAM surrounding the first exon of *GAS7* variant-c tended to be inversely correlated with mRNA levels (Figure 2).

**Table 5:**
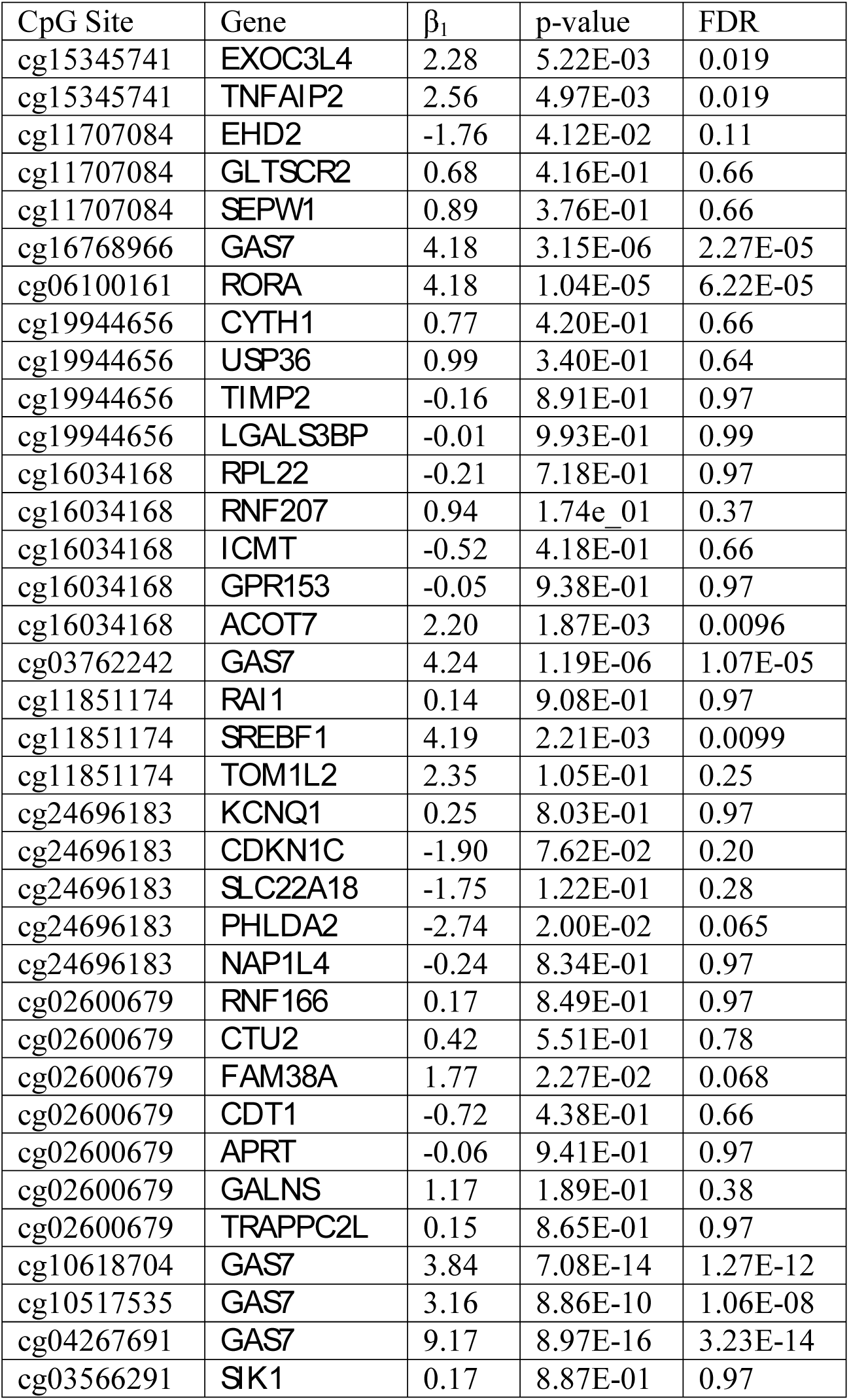
Expression quantitative trait methylation (eQTM) results for CpGs with PCT-adjusted meta-analysis p-values < 1e-05; analysis conducted with RICHS placental samples for which RNAseq and Illumina Infinium 450K were performed (n=200).

We then tested whether the expression levels of the FDR-significant genes from the eQTM analysis were associated with fetal growth metrics standardized by sex and gestational age (22) (Table 6). We found that the higher placental expression levels of *TNFAIP2* and *ACOT7* were modestly associated with decreasing birth weight z-scores, while higher expression of *RORA* was modestly associated with increased birth weight z-scores (p-values < 0.05). Higher expression of *ACOT7* was also associated with decreasing birth length and head circumference z-scores (p-values < 0.05). None of the other gene expression levels were associated with birth metrics.

**Table 6:** Associations between expression levels with growth metrics at birth in RICHS (n=200); birth weight (BW), length (BL), and head circumference (HC) were sex-specific z-scores that have been adjusted for gestational age, Tau represents the Kendall correlation coefficient.

|  | BW | BL | HC |
| --- | --- | --- | --- |
| Gene | Tau<br>(p-value) | Tau<br>(p-value) | Tau<br>(p-value) |
| *TNFAIP2* | -0.099<br>(0.039) | -0.057<br>(0.24) | -0.092<br>(0.061) |
| *EXOC3L4* | -0.024<br>(0.62) | 0.005<br>(0.91) | -0.030<br>(0.54) |
| *GAS7* | 0.012<br>(0.80) | 0.032<br>(0.51) | 0.032<br>(0.51) |
| *RORA* | 0.106<br>(0.026) | 0.068<br>(0.16) | 0.077<br>(0.12) |
| *ACOT7* | -0.134<br>(0.0048) | -0.106<br>(0.029) | -0.145<br>(0.0032) |
| *SREBF1* | 0.032<br>(0.50) | 0.003<br>(0.95) | -0.051<br>(0.30) |

Finally, we explored whether top hits from a previous small sample study (n=24) of placental Cd and DNA methylation levels (19) demonstrated associations in our study. In our meta-analysis one of the top six loci from that study, cg04528060 on chromosome 4 within the ADP-ribosylation factor-like 9 (*ARL9*) gene, demonstrated a nominally significant association with Cd (β_1_ = -0.49, p-value = 0.014) with the same direction of effect in our meta-analysis. However, we did not observe any evidence of sex-specific effects at this, or the other 5 CpG sites. We also explored whether any of CpGs from our meta-analysis with p-values < 1e-05 yielded statistically significant interactions between Cd and fetal sex (Supplemental Table 7). Of the 17 models, only cg02600679 within the gene body of *FAM38A* produced a significant interaction (p-value = 0.031). Stratifying by fetal sex revealed that Cd was associated with increased DNAM at cg02600679 among males (β_1_ = 0.26, p-value = 0.0012), while there was no association among females (p-value = 0.97).

## Discussion

We conducted a large EWAS study across two cohorts to examine associations between Cd concentrations and DNAM from human placentae and considered the confounding effects of placental tissue heterogeneity. We further assessed eQTM and associations with birth metrics. We used identical measurement techniques for both DNAM and trace metal measurement in the two cohorts, and the placental concentrations of Cd were similar to the 3.53 ng/g previously reported in the full NHBCS cohort (23), and towards the lower end but consistent with other cohorts that have reported concentrations ranging as low as 1.2 ng/g to as high as 53.3 ng/g (24). We identified Cd-associated variations at two CpGs within a 5% FDR threshold and an additional 15 CpGs with meta-analysis p-values < 1e-05. At all but one of these loci, higher Cd was associated with increased methylation levels. We also found that the methylation levels at these CpGs were associated with the expression patterns of six genes, and the expression of three genes (*TNFAIP2*, *ACOT7*, and *RORA*) were associated with birth metrics.

The observed preference for Cd-associated hypermethylation, as opposed to hypomethylation, is consistent with previous studies that observed higher Cd to be associated with a higher proportion of hypermethylation at gene promoters from fetal and maternal blood samples (25) and with increased methylation at regulatory regions for imprinted genes from cord blood samples (26). In contrast, others have observed Cd-associated hypomethylation in placental DNA, however this was from a relatively small study that focused on sex-specific associations (19). Cd has been shown to enhance DNA methyltransferase activity and thus increase the preponderance of hypermethylated loci, though these effects on DNAM may vary by timing and duration of exposure (27).

The top hit from our study, cg15345741 is on chromosome 14, approximately 70kb and 97kb downstream of the *TNFAIP2* and *EXOC3L4* genes, respectively, which encode the *tumor necrosis factor, alpha-induced protein 2* and the *exocyst complex component 3 like 4* protein. Higher methylation at cg15345741 was associated with higher expression of both *TNFAIP2* and *EXOC3L4*, and thus may be involved in the regulation of these genes or as a possible marker of their expression, though it does not function through the promoter methylation paradigm. However, the location of this CpG does overlap with a DNase hypersensitivity region and multiple putative transcription factor binding sites. We also identified the Cd-associated loci cg11707084 on chromosome 19 which is cis to six genes (*SEPW1*, *GLTSCR2*, *EHD2*, *TRPX1*, *CRX*, and *SULT2A1*), though the expression levels of these genes were not detectable in placental tissue or those that were did not yield associations with DNAM at our candidate loci. Thus, the potential functional role of DNAM at cg11707084 is unclear.

Our top 250 sites included genes that were significantly enriched for nitric oxide (NO) and G-protein signaling pathways (ras, apelin, oxytocin, and cGMP-PKG), cancerous process, cellular adhesion, cellular metabolism, and cytotoxicity. These pathways are consistent with currently recognized mechanisms of Cd-associated reproductive toxicity that have primarily been studied in animal or *in vitro* models: altered signal transduction, impaired cellular adhesion, disruptions to cell-cycle, increased oxidative stress and cytotoxic/apoptotic signaling (3). We provide evidence that these biologic processes in the human placenta may be perturbed by Cd-associated differential methylation, despite overall Cd-exposures being low and our study samples primarily consisting of healthy pregnancies. Furthermore, the functions of the genes whose expression levels were associated with our top hits are involved in similar mechanisms.

The proteins produced by both *TNFAIP2* and *EXOC3L4* are structurally similar to subunits of the exocyst, a molecular complex that orchestrates numerous important developmental functions including cellular membrane outgrowth, establishing cell polarity, and mediating cell-to-cell adhesion and communication (28). Given the crucial roles that the above processes play during development, it is not surprising that the exocyst is critical to proper placental development. Mouse models have demonstrated that full knockout of the exocyst subunits can cause early death of the embryo (28), while disrupted exocyst function may be associated with preeclampsia (29). Both *TNFAIP2* and *EXOC3L4* also serve other biological roles that may be important to Cd-associated toxicity. For instance, the expression of *TNFAIP2* can be up-regulated by TNFα, retinoic acid, interleukin-1β, and other pro-inflammatory and cytotoxic signals (30, 31). It is also highly expressed in some cancerous tissues, is related to poor survival, and is involved in cellular motility, invasion, and metastasis (32). Additionally, *TNFAIP2*, among other pro-inflammatory genes, has also been shown to be overexpressed in the hippocampus of Alzheimer’s disease patients (33) and may mediate cell death in motor neurons of amyotrophic lateral sclerosis patients (34), suggesting a role in neurodegenerative disease. SNPs within *EXOC3L4* have been implicated to affect gamma-glutamyl transferase (GGT) activity (35), and elevated levels of GGT are recognized markers of oxidative stress (36), which plays a central role in Cd-toxicity. Disrupted epigenetic regulation of these genes and their involvement in placental growth, development, inflammation and/or oxidative stress responses present compelling potential mechanisms through which Cd may elicit some of its reproductive toxicity.

Expression of the *GAS7* gene, which encodes the *growth arrest specific 7* protein, was strongly associated with DNAM at all five candidate CpGs that were associated with Cd (p-values < 1e-05). Interestingly, the strongest Cd-associations were localized nearby the first exons of transcript variants a, b, and d for *GAS7*, and thus these variations may play roles in Cd-associated alternative splicing of this gene. However, read-depth of our RNA-seq data was not sufficient to accurately test for variant-specific expression levels, and this should be investigated further in future studies given the potential for variant-specific functions described below. The *GAS7* gene, which is highly expressed in embryonic cells (37), mature brain cells and Purkinje neurons, plays an essential role in neurite outgrowth (38) and inducing neuronal cell death (39), specific functions may be isoform dependent. In the mouse embryo, levels of *GAS7* decrease as the embryo grows, except for in neuronal-specific epidermal cells in which expression remains high (37). *GAS7* expression also plays an essential role in other non-neuronal growth functions, such as osteoblast differentiation and bone development (40). Furthermore, neural outgrowth and arborization are shared functions between the exocyst and *GAS7* (28, 38), while *TNFAIP2* expression has been associated with some neurodegenerative conditions (33, 34). Relatively few studies have investigated the relationship between perinatal Cd exposure and neurodevelopment, though some epidemiologic studies suggest a relationship with impaired cognition in childhood (41). This raises some questions about the possible roles of the above genes in Cd-associated cognitive and neurobehavioral deficits. However, the functional roles of these genes in human placental tissue are not well understood. Thus it is unclear how, or whether, Cd-associated differential placental DNAM and expression may affect early life cognitive function and should be the focus of additional research.

Genes known to contribute in neurite outgrowth are sometimes also involved in regulating placental angiogenesis, as has been shown for vascular endothelial growth factor (VEGF) and neurotrophins (42). Interestingly, sterol regulatory element binding proteins (SREBPs), such as *SREBF1* whose expression was strongly correlated with one of our top hits, play key roles in VEGF-induced angiogenesis (43). Additionally, inappropriate placental vascularization can result in fetal growth restriction or other pregnancy complications (42). In our study, the expression of *GAS7*, *EXOC3L4*, and *SREBF1* were not associated with standardized percentiles for birthweight, head circumference, or birth length, and thus may not be directly related to fetal growth. Whereas expression of *TNFAIP2*, *ACOT7*, and *RORA* were associated with fetal growth. *ACOT7,* which encodes *acyl-CoA thioesterase 7*, is involved in lipid metabolism in neuronal cells and may prevent neurotoxicity (44), while it also promotes pro-inflammatory activities by up-regulating arachidonic acid production (45). *RORA*, which encodes the *retinoic acid receptor-related orphan receptor alpha*, is involved in numerous developmental processes including innate immune system development, regulating inflammatory responses, neuronal survival and growth, as well as lipid and glucose metabolism (46). One common biological thread that differentiates *TNFAIP2*, *ACOT7*, and *RORA*, from *GAS7*, *EXOC3L4*, and *SREBF1*, is their role in mediating inflammatory and immune responses. Thus, although we observed numerous Cd-associated variations in DNAM among genes involved in neurodevelopment and cellular growth processes, variations involved in inflammatory and immune signaling disruption may impact fetal growth.

Other epidemiologic studies of maternal Cd and birth/pregnancy outcomes, that also investigated molecular mechanisms, have found that higher Cd may affect essential metals transport to the fetus (13), reduce expression of a protocadherin, which are involved in neuronal development (47) and affect the DNAM of genes involved in cell death, lipid metabolism, and cancer pathways (25). Similarly, electronic waste (e-waste) exposure, which is associated with higher placental concentrations of toxic metals including Cd, has been associated with differential expression of proteins primarily involved in immune and metabolic processes (48). The top hits from a small sample EWAS of placental Cd and DNAM observed differential methylation of genes involved in cellular metabolism, growth, and cell damage response (19). We were able to reproduce Cd-associated DNAM variations for one top hit from this study, cg04528060 in the 5’UTR of the *ARL9* gene, (meta-analysis P-value < 0.05) with the same direction of effect. Interestingly, the *ARL9* gene encodes for a GTP-binding protein that is structurally similar to members of the RAS superfamily (49), and RAS-signaling was the most highly enriched KEGG pathway from our study.

The preponderance of similar biological processes associated with Cd across multiple studies with different technologies, populations, and tissues, is striking, and suggests that common mechanisms may be affected by Cd exposure during pregnancy. These findings should be interpreted within the context of this study’s limitations. This was an observational study in which placental Cd concentrations and DNA-M levels were both measured in placenta at term. Thus, we cannot rule out the possibility of reverse causation since exposure and outcome were measured at the same time-point, and the observed associations at term may not be representative of Cd and DNA-M associations throughout development. Though we adjusted for likely confounders in our study, we also cannot rule out the possibility that unmeasured or residual confounding may have affected our observations. However, to our knowledge this was the largest study yet to examine the relationships between placental Cd and DNAM, and the first account for and evaluate associations with estimated cellular heterogeneity.

The loci that we did identify are involved in biological processes known to be affected by Cd-toxicity and thus are strong candidates for future studies. We observed these associations in two healthy populations with relatively low exposure levels. In the US, exposure to Cd has declined since the late 1980s, which is primarily attributed to declining smoking rates (50). However, Cd is still detectable in numerous commonly consumed foods (51) and exposure prevalence remains quite high (52). Additionally, e-waste recycling and disposal are emerging sources of environmental Cd-contamination (and other toxic metals), particularly in developing countries (53). This raises some concerns about the potential threshold for Cd-associated reproductive toxicity, and whether current dietary intake guidelines and efforts to reduce tobacco smoke exposure during pregnancy adequately protect against adverse birth and pregnancy outcomes. We encourage further molecular epidemiologic studies of the associations between Cd and DNAM in placental tissue, particularly within populations with higher exposure levels and different ethnic backgrounds, but also encourage studies of the associations between dietary intake and smoking-related Cd exposure with the placental accumulation of Cd. Understanding these relationships can help us determine whether changes to current guidelines and recommendations could reduce Cd-associated adverse reproductive outcomes. Finally, the genes whose expression levels were associated with DNAM at our top hits are multifunctional, and involved in neurodevelopmental and/or neurodegenerative processes, cellular growth processes, and inflammatory/immune signaling. We also found that the expression levels of three of these genes were associated with infant birth weight. Thus, additional future studies should investigate whether differential placental epigenetic regulation and expression of *GAS7*, *EXOC3L4*, *SREBF1*, *RORA*, *ACOT7* and *TNFAIP2*, are associated with cognitive/neurobehavioral, growth, and immune outcomes in children.

## Methods

### New Hampshire Birth Cohort Study

This study included mother-infant pairs from the NHBCS, an ongoing birth cohort initiated in 2009. Women that were currently pregnant, between 18 and 45 years of age, receiving prenatal care from one of the study clinics in New Hampshire, reporting that the primary source of drinking water at their residence was from an unregulated well, and having resided in the same household since their previous menstrual period with no plans to move before delivery, were enrolled in the cohort. All participants provided written informed consent in accordance with the requirements of the Institutional Review Board (IRB) of Dartmouth College. The sample for this study consisted of NHBCS participants that were recruited between February 2012 and September 2013, for whom placenta were sampled to conduct genetic and epigenetic assays (n=343). Placental gross measures such as placental diameter (cm) and placental weight (g) were collected immediately after delivery. Interviewer administered questionnaires and medical record abstraction were utilized to collect sociodemographic, lifestyle, and anthropometric data.

### Rhode Island Child Health Study

The RICHS enrolled mother-infant pairs with non-pathologic pregnancies at the Women and Infants’ Hospital in Providence, RI, USA between September 2010 and February 2013. Exclusion criteria consisted of mothers younger than 18 years of age, with life threatening conditions, pregnancies with gestational time < 37 weeks, or infants with congenital/chromosomal abnormalities. All protocols were approved by the institutional review boards at the Women and Infants Hospital of Rhode Island and Dartmouth College and all participants provided written informed consent. Infants that were born small for gestational age (≤ 10^th^ BW percentile) or large for gestational age (≥ 90^th^ BW percentile) were oversampled, then infants adequate for gestational age (between the 10^th^ and 90^th^ BW percentiles) that matched on gestational age and maternal age were coincidentally enrolled. This study included all mother-infant-pairs for whom placental metal concentrations and placental DNA-M arrays had been conducted (n=141). Interviewer administered questionnaires were utilized to collect sociodemographic and lifestyle data; anthropometric and medical history data were obtained via structured medical records review.

### Sample Collection

Placentae samples from RICHS and NHBCS were biopsied from the fetal side adjacent to the cord insertion site, within 2 hours of delivery and maternal decidua was removed. Samples were placed in RNAlater (Life Tecnologies, Carlsbad, CA) then frozen at -80°C. Both RNA and DNA were extracted (Norgen Biotek, Thorold, ON) then quantified via the Qubit Flourometer (Life Technologies), and subsequently stored at -80°C.

### DNA Methylation, QC, and Normalization

Both cohorts measured DNAM via Illumina Infinium HumanMethylation450K BeadArray (Illumina, San Diego, CA) at the University of Minnesota Genomics Center. Bisulfite modification was done with the EZ Methylation kit (Zymo Research, Irvine, CA), samples were randomized across multiple batches, and data was assembled using BeadStudio (Illumina). The raw array data are available via the NCBI Gene Expression Omnibus (GEO) for NHBCS and RICHS via accession numbers GSE71678 and GSE75248, respectively. For QC, we excluded probes with poor detection p-values (p-value > 0.001), measuring DNA-M at X- and Y-linked CpG sites, with a single nucleotide polymorphism (SNP) within 10bp of the target CpG or single base extension (SBE) (and allelic frequency > 1%), or cross-hybridizing to multiple genomic regions (54). Background correction, dye bias, and functional normalization were performed via the minfi package (55) and standardization across probe-types was performed via beta mixture quantile (BMIQ) normalization (56). Batch effects were corrected for using empirical Bayes via the combat function in R (57). Methylation data were reported as β-values, representing the proportion of methylated alleles for each individual CpG site. For the EWAS models described below, we utilized M-values, logit-transformation of the beta values, which better approximate a normal distribution (58).

### RNA sequencing, QC, and Normalization

Transcriptome-wide sequencing was performed on 200 placental samples from RICHS. We used RNeasy Mini Kit (Qiagen, Valencia, CA) to isolate total RNA, then stored at -80°C until analysis. We then quantified RNA via Nanodrop Spectrophotometer (Thermo Scientific, Waltham, MA), assessed integrity via the Agilent Bioanalyzer (Agilen, Santa Clara, CA), removed ribosomal RNA via Ribo-Zero Kit (59), converted to cDNA using random hexamers (Thermo Scientific, Waltham, MA), and performed transcriptome-wide RNA sequencing via the HiSeq 2500 platform (Illumina, San Diego, CA) (60). Raw reads have been deposited at the NCBI sequence read archive (SRP095910). Quality control was performed in FastQC, then reads were mapped to the human reference genome (h19) using the Spliced Transcripts Alignment to a Reference (STAR) aligner. Low-expressed genes were excluded, read counts were adjusted for GC content (61), then normalized via the trimmed mean of m-values (TMM) (62); final data are normalized log2 counts per million (logCPM) reads.

### Cadmium Quantification

Trace element concentrations were quantified in RICHS and NHBCS placental samples at the Dartmouth Trace Elements Analysis Core using inductively coupled plasma mass spectrometry (ICP-MS); details of the processing are described elsewhere (23). None of the RICHS samples received non-detectable Cd concentrations, while only four of the NHBCS samples received non-detects which were assigned a value equal to the 0.5 percentile of detectable values. For EWAS models, we dichotomized cadmium concentrations by the cohort-specific median concentrations (NHBCS median = 3.13 ng/g) and (RICHS median = 4.34 ng/g) for the EWAS analysis.

### Estimation of placental tissue heterogeneity

We addressed the potential issue of confounding due methylation-associated tissue heterogeneity via the RefFreeEWAS package in R (21). This method utilized non-negative matrix factorization to estimate the proportions of putative cellular mixtures and has been demonstrated to yield reliable estimates of the constituent cell types and the underlying methylomes that define them (21). Utilizing the 10,000 CpGs with the greatest variability in methylation and we identified four and five putative constituent cell mixtures in the RICHS and NHBCS cohorts respectively, then used the full set of CpGs for each cohort (NHBCS p=408,367 and RICHS p=397,040) to estimate the relative proportions of each putative cell-type per placental sample. We examined whether the 10,000 CpGs with the largest variance across putative cell-type specific methylation values were enriched for gene ontology (GO) terms or Kyoto Encyclopedia of Genes and Genomes (KEGG) pathways, via the gometh function in the missMethyl package in R (63). This method adjusts for the potential bias introduced by some genes having greater probability of being included in the gene-set due to having greater numbers of CpGs on the 450K array. We examined the consistency of the CpGs with high cell-type specific methylation, as well as consistency in the top 100 GO-terms and KEGG pathways associated with those highly variable CpGs.

### Epigenome-wide association study (EWAS)

Robust linear regressions (rlm) were used to estimate the associations between CpG-specific DNA methylation levels and high-vs-low placental concentrations of Cd. We produced two covariate-adjusted models for each CpG within each cohort, first regressing M-values on dichotomized Cd-concentrations (cohort-specific median as the reference) while adjusted for infant sex, maternal smoking during pregnancy, and highest achieved maternal education level. Second, we produced the same models while also adjusted for the estimated proportions of putative placental cell-types. Genomic inflation was examined via the genomic inflation factor (λ) which was calculated with the genABEL package in R (64). We selected the 250 CpGs with the smallest Cd-associated meta-analysis p-values and assessed these for functional enrichment with GO-terms and KEGG pathways as described above. Statistical significance for enrichment was determined at a 5% FDR.

### Meta Analyses

We used METAL to meta-analyze the cohort-specific Cd-associations via inverse variance weighted fixed-effects models, and evaluated interstudy heterogeneity via Cochran’s Q-test p-values < 0.05. (65). We produced FDR-adjusted p-values via the qvalue package in R. Statistical significance was determined at an FDR of 5%. Volcano plots were produced to visualize overall magnitude and direction of associations, while Manhattan plots were produced via the qqman package to visualize the genomic distribution of significant associations. Plots for specific regions of the genome were visualized using the coMET package (66). Top hits from the study were also evaluated for dose-response relationships via modeling associations between log-transformed continuous Cd and DNAM, via linear mixed models, allowing for random intercepts by study.

### Expression Quantitative Trait Methylation (eQTM) Analysis

We then tested whether any of our top Cd-associated CpGs could be expression quantitative trait methylation (eQTM) loci within the 200 RICHS samples, 85 of which overlap with the samples utilized in the EWAS. We tested for cis-gene-CpG associations (within 100kb of the target CpG site) by regressing gene expression levels (log_2_(CPM)) on CpG β-values using robust linear models. Statistical significance for eQTM associations was determined at a 5% FDR.

### Associations with fetal growth

We tested for associations between birth metrics and gene expression using Kendal correlations, which is a non-parametric correlation test that allows for ties. Since birth metric z-scores were standardized by fetal sex and by weeks of gestation, no additional adjustments were made to these association tests.

### Replication of previous studies and sex-specific effects

We then investigated whether we could reproduce the results from a previous study of sex-specific associations between placental Cd and placental DNAM (19). Because multiple studies of various fetal and maternal tissues have suggested that associations between Cd and DNAM may differ by fetal sex (7, 19, 67), we also screened the CpGs from our study with meta-analysis p-values < 1e-05 for interactions with fetal sex. For these analyses, we utilized linear mixed models with DNAM M-values as the outcome, allowing for random intercepts by study, and included an interaction-term between DNAM levels and fetal sex while adjusting for maternal education and maternal smoking. We utilized the Wald test to produce p-values, and determined statistically significant interactions at a p-value threshold of 0.05. Any models producing interaction p-values < 0.05 were then re-run, stratified by fetal sex.

## Acknowledgements

This work was supported by the National Institutes of Health [NIH-NIMH R01MH094609, NIH-NIEHS R01ES022223, NIH-NIGMS P20 GM104416 and NIH-NIEHS P01 ES022832] and by the United States Environmental Protection Agency [US EPA grant RD83544201]. Its contents are solely the responsibility of the grantee and do not necessarily represent the official views of the US EPA. Further, the US EPA does not endorse the purchase of any commercial products or services mentioned in the presentation.

